# High-throughput imaging of mRNA at the single-cell level in human primary immune cells

**DOI:** 10.1101/2020.11.10.377283

**Authors:** Manasi Gadkari, Jing Sun, Adrian Carcamo, Hugh Alessi, Zonghui Hu, Iain D.C. Fraser, Gianluca Pegoraro, Luis M. Franco

## Abstract

Measurement of gene expression at the single-cell level has led to important advances in the study of transcriptional regulation programs in healthy and disease states. In particular, single-cell gene expression approaches have shed light on the high level of transcriptional heterogeneity of individual cells, both at baseline and in response to experimental or environmental perturbations. We have developed a method for High-Content Imaging (HCI)-based quantification of transcript abundance at the single-cell level in primary human immune cells and have validated its performance under multiple experimental conditions to demonstrate its general applicability. This method, which we abbreviate as hcHCR, combines the high sensitivity of the hybridization chain reaction (HCR) for the visualization of mRNA molecules in single cells, with the speed, scalability, and technical reproducibility of HCI. We first tested eight microscopy-compatible attachment substrates for short-term culture of primary human B cells, T cells, monocytes, or neutrophils. We then miniaturized HCR in a 384-well format and documented the ability of the method to detect increased or decreased transcript abundance at the single-cell level in thousands of cells for each experimental condition by HCI. Furthermore, we demonstrated the feasibility of multiplexing gene expression measurements by simultaneously assaying the abundance of two transcripts per cell, both at baseline and in response to an experimental stimulus. Finally, we tested the robustness of the assay to technical and biological variation. We anticipate that hcHCR will be a suitable and cost-effective assay for low- to medium-throughput chemical, genetic or functional genomic screens in primary human cells, with the possibility of performing personalized screens or screens on cells obtained from patients with a specific disease.

## Introduction

Gene expression assays are the cornerstone of functional genomics. Measurement of transcript abundance is central to our current understanding of cell biology, defining the basal state of different cell types as well as their response to environmental or experimental perturbations. In recent years, the emphasis has been on the analysis of gene expression at the single-cell level. In the case of immune cells, this has shed light on the heterogeneity of the transcriptional state of individual cells and to the identification of transcriptional subsets among what were previously thought to be homogeneous populations (Stubbington et al. 2017). It has also advanced our understanding of the sets of functionally related and co-regulated genes that govern the response to stimuli by individual cells (Pope and Medzhitov 2018). Immune cell heterogeneity and transcriptional regulation at the level of individual cells are especially relevant at a time when the development of cancer immunotherapy has accelerated (Gibellini et al. 2020), the effects of immunosuppressive drugs are being revealed to be highly cell type-dependent (Franco et al. 2019), and a rapidly growing arsenal of new drugs targeting specific components of immune signaling networks is being developed (Tiligada et al. 2015).

The improved ability to study gene expression at the level of individual cells has been driven by a series of technological advances that have moved the experimental toolkit in two opposite but complementary directions: one with greater breadth and the other with greater depth. Greater breadth has been achieved by advances in single-cell transcriptomics. Single-cell RNA sequencing (scRNA-seq) allows simultaneous gene expression measurements of hundreds to thousands of genes per cell (Hwang et al. 2018) and has become the method of choice for identifying transcriptional subsets of cells. The high cost per sample and limited scalability have so far limited the use of scRNA-seq to low-throughput applications. In addition, current scRNA-seq technologies rely on a superficial sampling of each cell’s transcriptome, making the sensitivity of detection of a transcript in any given cell low and biasing the representation towards more highly expressed genes (Chen et al. 2019). Greater depth has been achieved by concomitant advances in RNA fluorescence in situ hybridization (FISH) methods, which have greatly increased the sensitivity of detection of individual RNA molecules (Pichon et al. 2018). This higher sensitivity enables the reliable detection of transcripts with low expression levels and the quantification of small changes in transcript abundance in large populations of cells under thousands of experimental conditions, but it comes at the expense of multiplexing, as the number of transcripts that can be assayed simultaneously is limited by the number of available fluorescence channels, usually up to 4 or 5 for advanced microscopy equipment.

High-content imaging (HCI) employs automated liquid handling, image acquisition, and image analysis, and offers the possibility of quantitative analyses at the single-cell level with high throughput and limited inter-operator variation (Pegoraro and Misteli 2017; Esner et al. 2018). The possibility of testing hundreds to tens of thousands of experimental conditions makes HCI particularly suitable for medium- or high-throughput screening experiments aimed at understanding the cellular effects of large collections of perturbing agents like chemical compounds, RNAi, or CRISPR/Cas9. Such screening experiments are a central component of drug development pipelines (Hughes et al. 2011) and are a powerful tool for dissecting signaling networks in biological systems by selective manipulation of individual components (Sun et al. 2016). Most HCI assays to date have relied on the detection of signals from fluorescent dyes, stably expressed fluorescent proteins, or fluorescently labeled antibodies directed against endogenous proteins of interest. However, the same principles can be applied to high-throughput quantitative measurements of gene expression (Querido et al. 2017).

HCI assays have also relied primarily on immortalized and/or transformed cancer cell lines, which are easier to grow and manipulate than primary cells, but also tend to have substantial structural genomic abnormalities, which can limit the generalization of the results of transcript-level assays obtained in these cells to more physiologically relevant systems (Mittelman and Wilson 2013; Gioia et al. 2018; Zhou et al. 2019). In addition, cell lines can have different responses to chemical stimuli when compared to primary cells or cells exposed in vivo (Lavrentieva 2018). Finally, human cell lines are generally derived from a single individual, which limits the generalizability of the results as they cannot account for biological, inter-individual variation. Screening assays based on primary human cells would overcome these issues and would also allow for personalized screens with cells obtained directly from specific patients, or from patients with a particular disease of interest (Lavrentieva 2018). However, their culture is more technically challenging due to cell-to-cell heterogeneity, variable attachment properties, and low proliferation capacity (Hauser 2015; Lavrentieva 2018).

To address these limitations, we have developed a high-throughput method for quantitative analysis of gene expression at the single-cell level in primary human cells. This method, which we abbreviate as hcHCR, combines HCI with a recently developed chemistry for high-sensitivity RNA FISH, the hybridization chain reaction (HCR) (Choi et al. 2010, 2018). To demonstrate the general applicability of hcHCR, we have validated its performance on technical and biological replicate experiments with different primary human cell types and under multiple experimental conditions.

## Results

### High-content imaging-based quantification of transcript abundance at the single-cell level

The assay workflow for hcHCR is summarized in Fig. 1. Primary cells from human peripheral blood are first isolated and plated with cell culture media in 384-well imaging plates (Fig. 1A). After a rest period, intended to allow stabilization of gene expression after plating, the cells are treated, simultaneously or in tandem, with one or more chemical or biological perturbing agents whose effect on gene expression are being tested. Treated cells are then fixed and RNA is hybridized in situ with sequence-specific oligo DNA probe sets carrying HCR initiator sequences, followed by HCR amplification and automated image acquisition (Fig. 1A) (Choi et al. 2010, 2018). High-Content Image analysis is then used to first segment nuclei based on the DAPI image, followed by dilation of a mask to cover the cell body (cell segmentation), and by HCR spot detection and counting (Fig. 1B). The relative abundance of up to 3 specific mRNA transcripts can be quantified at the single-cell level in up to thousands of wells on a commercial HCI instrument. Gene expression can be quantified as spot counts per cell (digital HCR; Choi et al. 2018, and Fig. 1C left panel). Alternatively, when very high transcript abundance renders the density of individual HCR spot signals too high to be optically resolved, gene expression can be quantified as HCR mean fluorescence over the cell body region (quantitative HCR; Choi et al. 2018, and Fig. 1C center panel). In multiplexed experiments, when more than one gene is being imaged in each cell, hcHCR image data can be converted to the FCS format for visualization and analysis with standard flow cytometry software, including gating and bivariate scatterplots (Fig. 1C, right).

**Figure 1.**
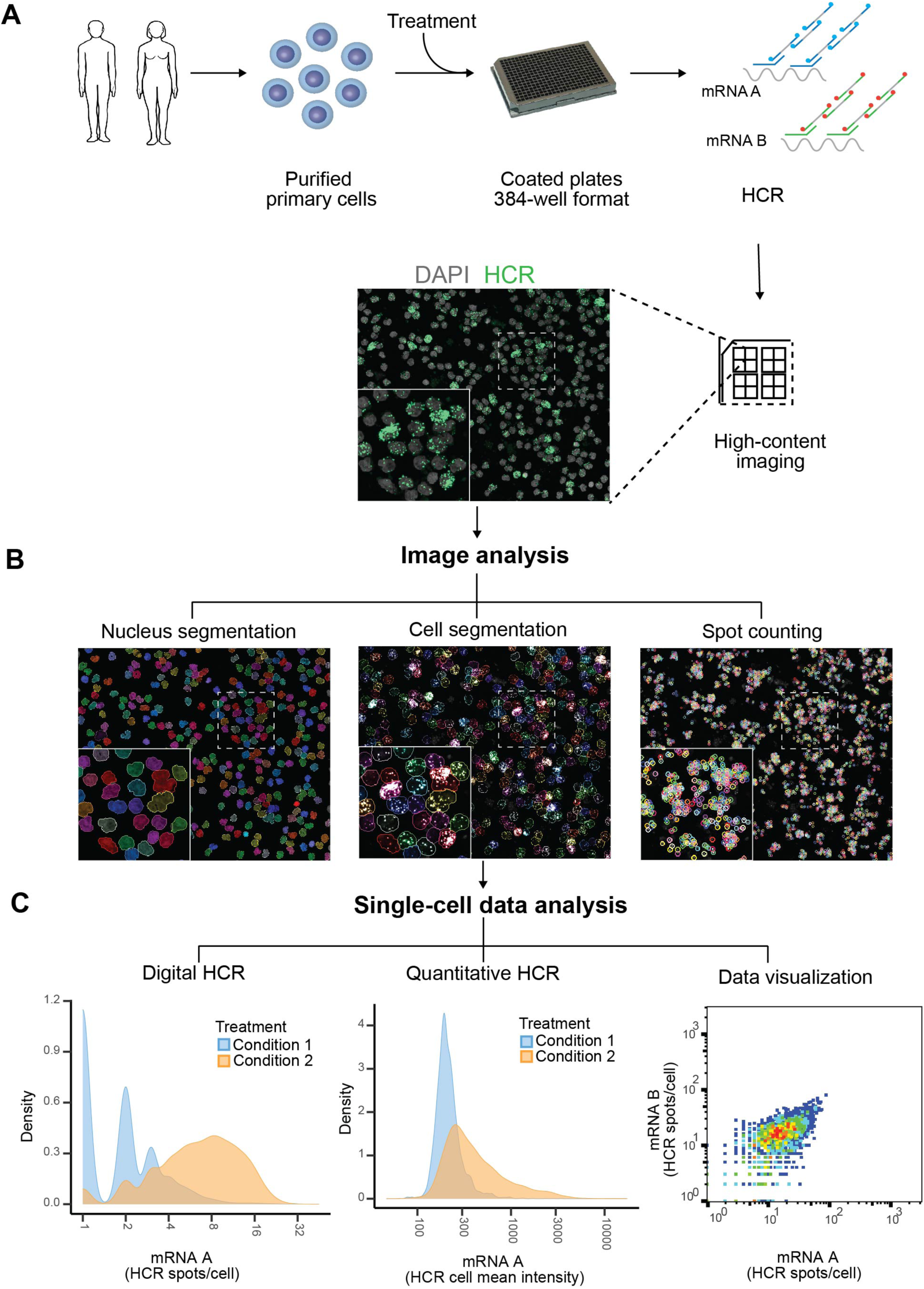
Schematic representation and workflow of the hcHCR method. (A) Primary human cells are purified and plated in 384-well imaging plates followed by in vitro treatment, RNA HCR, and high-content quantitative imaging at single-cell resolution. (B) High-Content Image analysis of HCR images: nuclei regions, cell body regions, and HCR spot locations are shown in pseudocolors. (C) Analysis of hcHCR data. Digital HCR (dHCR) involves mRNA quantification as HCR spots/cell. A representative single-cell density plot is shown on the left panel. Quantitative HCR involves quantification of HCR mean fluorescence intensity over the cell body region. A representative single-cell density plot is shown on the center panel. Digital or quantitative hcHCR data can be analyzed and gated with standard flow cytometry software. A bivariate scatter plot of dHCR counts for two genes is shown on the right panel.

### Identification of appropriate substrates for HCI of human primary immune cells

Fluorescence microscopy acquisition is greatly facilitated by the attachment of cells to the bottom of imaging plates. With this goal in mind, we began by comparing the adherence of four lymphoid or myeloid primary human immune cell types, which normally grow in suspension, to different substrates in 384-well imaging plates. Purified B cells, monocytes, neutrophils, or CD4+ T cells were plated live in wells coated with one of eight substrates: MS-1, MS-2, MS-3, 3D Hydrogel, PDL, SPA, 3D Hydrogel PDL, or 3D Hydrogel SPA. After a refractory period of 2 to 4 hours to allow for adherence to the substrate and stabilization of gene expression after plating, cells were fixed. Culture plates were then subject to the same incubation conditions and washes that would normally be used in our hcHCR protocol, but without the addition of HCR probes or hairpins (see Materials and Methods). Cell nuclei were then stained with DAPI and cell attachment was quantified by HCI (Fig. 2A).

**Figure 2.**
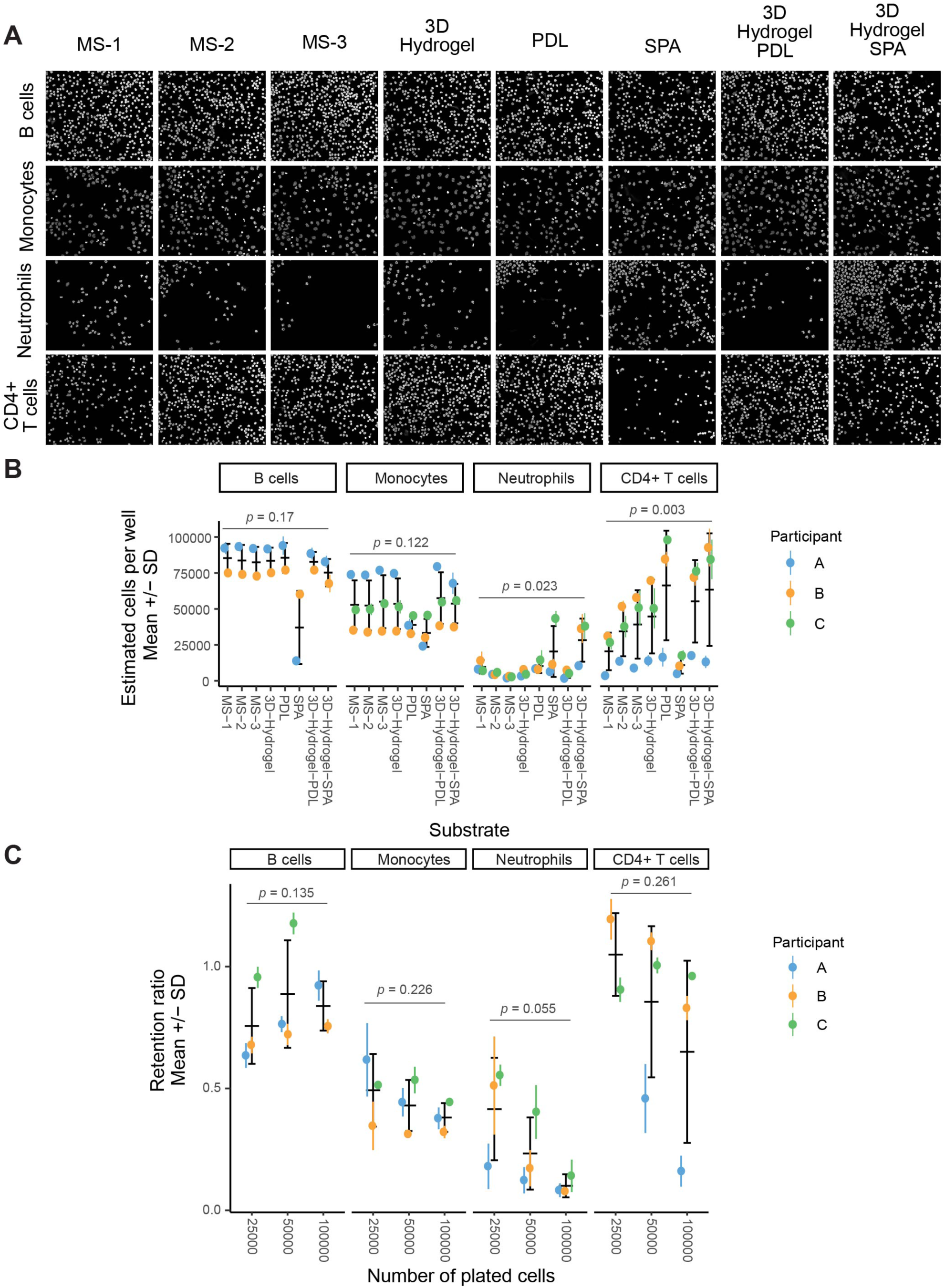
Identification of an appropriate substrate for HCI of human primary immune cells. Four primary human immune cell types (B cells, monocytes, neutrophils and CD4+ T cells) were independently cultured in 384-well plates in which well bottoms were coated with 1 of 8 substrates: MS-1, MS-2, MS-3, 3D Hydrogel, PDL, SPA, 3D Hydrogel PDL, or 3D Hydrogel SPA. Each cell type was plated at three concentrations: 100,000, 50,000 or 25,000 cells/well, in technical replicates. (A) Representative images of each cell type and substrate at a concentration of 100,000 cells/well. Cells were fixed with 4% PFA and stained with DAPI prior to imaging. (B) Cell attachment for 4 primary human immune cells in 8 substrates. Cells were plated at 100,000 cells/well. The y-axis represents the number of cells counted after fixation and automated liquid handling in conditions similar to those of the hcHCR protocol. Each dot represents one biological replicate (one unrelated healthy human donor). Colored error bars display the SD for 3 technical replicates of each biological replicate. Black error bars display the mean ± SD of the biological replicates. Significance values are from a linear mixed-effects model. (C) Retention ratios for 4 primary human immune cells at 3 cell concentrations. Cells were plated in PDL substrate and underwent fixation and automated liquid handling in conditions similar to those of the hcHCR protocol. The retention ratio for each biological replicate was calculated as the estimated number of cells per well [(number of cells counted)/ (number of fields of view imaged) × (total number of fields of view per well)] divided by the number of cells plated per well. Each dot represents one biological replicate (one unrelated healthy human donor). Colored error bars display the SD for 3 technical replicates of each biological replicate. Black error bars display the mean ± SD of the biological replicates. Significance values are from a linear mixed-effects model.

Cell attachment was comparable among the 8 substrates for B cells and monocytes (Fig. 2B). While the mean number of cells retained showed no statistically significant difference among the substrates, the mean number of cells retained was lower in SPA than in other substrates (Fig. 2B). In contrast, neutrophils were retained better in 3D hydrogel with SPA than in any of the other substrates tested (Fig. 2A and Fig. 2B). CD4+ T cells showed significant differences in attachment across substrates, with PDL and 3D hydrogel-based substrates having the highest attachment, and SPA again being the substrate with the lowest attachment. We chose PDL as a substrate for subsequent experiments, given its lower cost and greater availability.

Because the number of available primary human cells is often limited in practice, we then tested whether the retention ratios for each of the 4 cell types, cultured in PDL substrate, would change over a range of cell concentrations. We found no statistically significant differences in retention ratios for any of the 4 cell types tested, at concentrations of 25,000, 50,000, or 100,000 cells per well (Fig. 2C).

The results of these experiments indicate that different primary immune cell types can be successfully cultured short-term in 384-well imaging plates coated with a variety of substrates, with sufficient cell retention after fixation and automated liquid handling to allow for HCR followed by HCI.

### Detection of up- or down-regulation of gene expression in human primary immune cells

We then tested whether hcHCR could be used to quantify up-regulation or down-regulation of gene expression at the single-cell level in primary immune cells (Fig. 3). As an example of gene expression increase, we measured transcript abundance for the gene *TSC22D3 (GILZ)*, a classic glucocorticoid-inducible gene which has been studied extensively in human and animal models (D’Adamio et al. 1997; Cannarile et al. 2001). As expected, primary human B cells treated for 2 hours with the glucocorticoid methylprednisolone (MP) showed a four-fold up-regulation of *TSC22D3* (*GILZ*) compared to vehicle-treated cells, measured as the number of *TSC22D3* HCR spots per cell (Fig. 3A and 3B). As an example of gene expression decrease, we measured transcript abundance for the gene *TNF*, which encodes the inflammatory cytokine tumor necrosis factor alpha (TNF), in primary human monocytes treated with MP for 2 hours after 30 minutes of lipopolysaccharide (LPS) stimulation. In response to LPS, monocytes are known to produce large quantities of *TNF* by induction of gene expression (Chen et al. 1985; Kornbluth and Edgington 1986), whereas glucocorticoids like MP are known to suppress this induction (Hodge et al. 1999; Waage and Bakke 1988). Upon sequential LPS stimulation and MP treatment, and as compared to LPS stimulation alone, we were able to detect a four-fold decrease in *TNF* transcript abundance, measured as the number of *TNF* HCR spots per cell (Fig. 3C and 3D).

**Figure 3.**
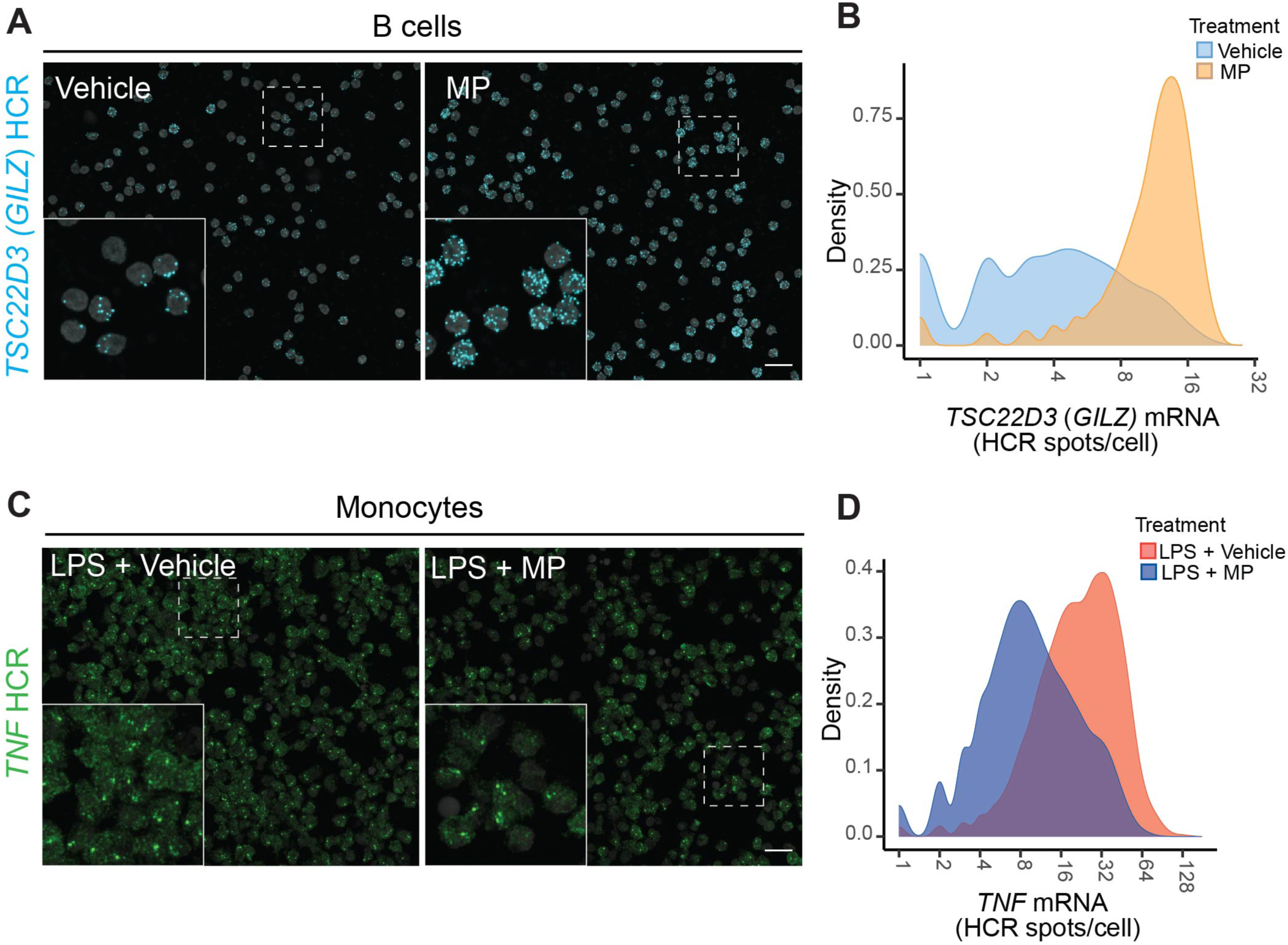
Detection of up- or down-regulated genes. (A) *TSC22D3* (*GILZ*) transcript abundance as HCR spots/cell (blue) after in vitro treatment of human primary B cells with vehicle (0.1% ethanol) or methylprednisolone (MP) (200 µg/dL) for 2 hours. (B) Density plots showing the distributions of *TSC22D3* transcript abundance after MP or vehicle treatment. (C) *TNF* transcript abundance as HCR spots/cell (green), after in vitro stimulation of human primary monocytes with LPS (1 ng/mL) for 30 minutes, followed by treatment with vehicle (0.1% ethanol) or MP (200 µg/dL) for 2 hours. (D) Density plots showing *TNF* mRNA quantification after LPS stimulation followed by vehicle or MP treatment.

These results indicate that hcHCR can reliably measure up-regulation or down-regulation of gene expression upon treatment of human primary immune cells with different chemicals.

### Simultaneous quantification of transcript abundance for multiple genes at the single-cell level

One of the most appealing properties of HCR is that this technique can simultaneously measure the expression of several RNA transcript species in the same cell (Choi, 2018). We sought to test multiplexed, high-throughput HCR by simultaneously incubating primary human monocytes with DNA oligo probe sets against *TNF* with one barcode, and against the gene encoding another inflammatory cytokine, interleukin 1 beta (*IL1B*) with a different barcode. Samples were then stained with barcode-specific HCR amplification hairpins labeled with different fluorophores. We performed a *TNF* and *IL1B* mRNA up-regulation time course by stimulating monocytes with LPS for increasing amounts of time, fixing them, then performing HCR staining as described above (Fig. 4A). As expected, an increasing number of HCR foci in both fluorescence channels appeared with longer exposure of monocytes to LPS (Fig. 4A), indicating up-regulation of *TNF* and *IL1B* mRNA expression in these cells. Single-cell analysis of *TNF* and *IL1B* mRNA expression by hcHCR 30 minutes after vehicle or LPS stimulation revealed a strong positive correlation between expression of *TNF* and *IL1B* (Fig. 4B). Visual inspection of the images generated in the presence of LPS at 15 minutes and subsequent timepoints revealed the appearance of one or two large bright nuclear *TNF* or *IL1B* HCR foci, which likely represent sites of active transcription at each of the two alleles of each gene (Fig. 4A and 4C). Expanding on these initial observations, we then used hcHCR to test a matrix of experimental conditions in which the LPS concentration and/or the length of LPS exposure were changed (Fig. 4D). Using this approach, we could document a robust increase of both *TNF* and *IL1B* expression in the population of monocytes after 60 minutes in the presence of LPS 0.01 ng/mL when compared to baseline (Fig 4D). Furthermore, at LPS concentrations of 0.1 ng/mL or higher, the induction of *TNF* and *IL1B* expression was already evident at 30 minutes (Fig 4D). Finally, at these concentrations, longer times of exposure to LPS (60 or 120 minutes) did not result in further measurable increases in *TNF* or *IL1B* expression as measured by hcHCR, possibly indicating a saturation of the of the dHCR signal or the induction of negative regulatory mechanisms (Fig 4D). The results of these experiments demonstrate that hcHCR can detect the expression of multiple mRNA transcripts and quantify co-expression at the single-cell level. They also indicate that hcHCR can rapidly and quantitatively test a range of different experimental conditions employing a limited number of human primary immune cells. Given that hcHCR detected robust upregulation of *TNF* and *IL1B* at 1 ng/mL LPS for 30 minutes, we decided to use these experimental conditions to conduct further experiments to test the performance of the assay across biological replicates.

**Figure 4.**
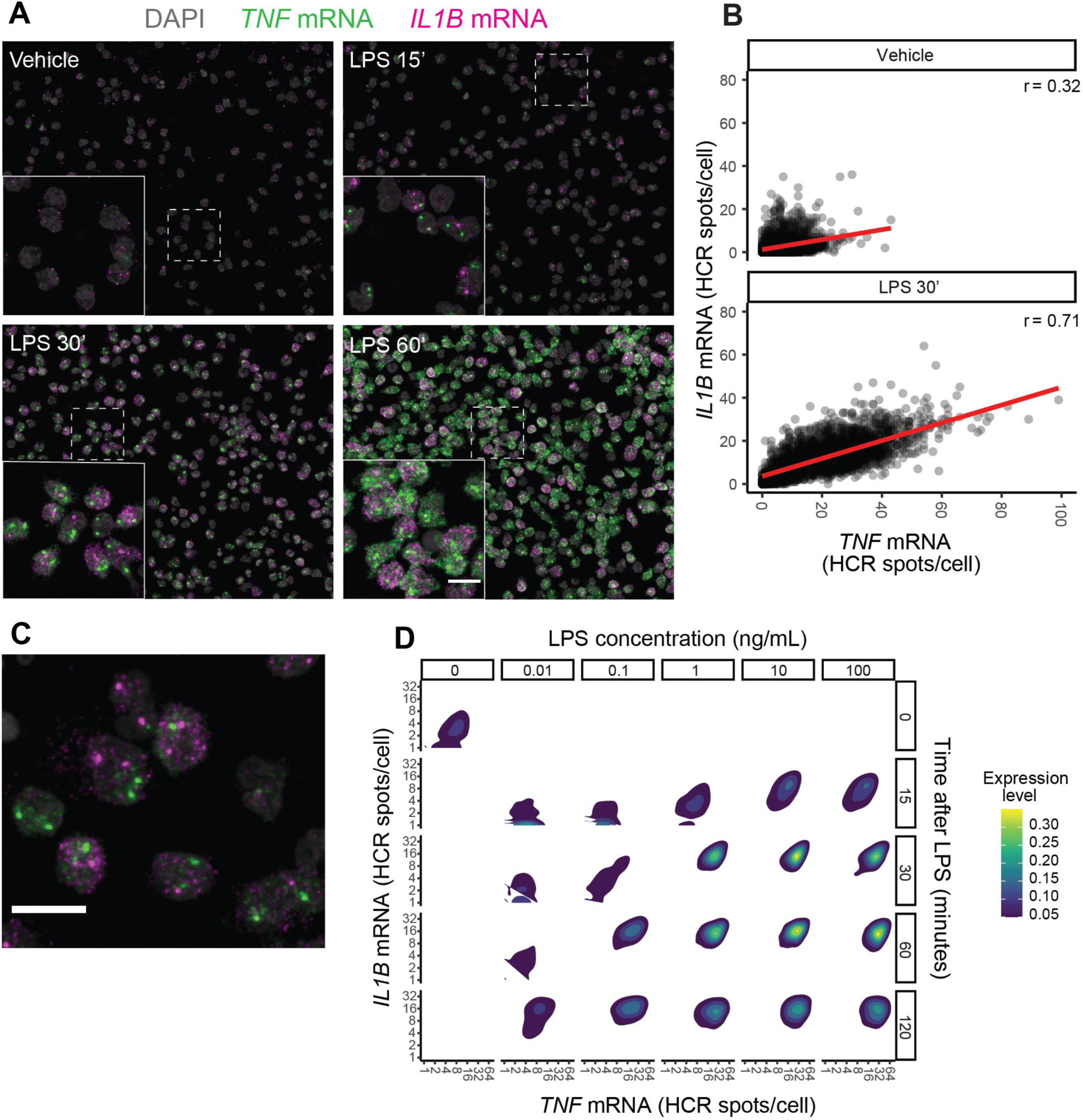
Simultaneous quantification of transcript abundance for multiple genes. Single-cell level quantification of gene expression for two genes, *TNF* (green) and *IL1B* (magenta), assayed in the same reaction in primary human monocytes stained with DAPI (grey). (A) Time-series response to stimulation with LPS (1 ng/mL), showing *TNF* and *IL1B* mRNA spots with vehicle or LPS stimulation at 15, 30 and 60-minute time points. (B) Scatter plots of *TNF* and *IL1B* gene expression, quantified as the number of HCR spots per cell. Cells were stimulated with vehicle or LPS for 30 minutes before hcHCR. Least-squares regression lines are in red. r = Pearson correlation coefficient. (C) Distinct clusters of mRNA molecules for each gene after stimulation with LPS (10 ng/mL) for 30 minutes. (D) Matrix of *IL1B* and *TNF* expression distributions after LPS stimulation at six concentrations (0, 0.01, 0.1, 1, 10, or 100 ng/mL) and five time points (0, 15, 30, 60, or 120 minutes).

### Quantitative assessment of technical and biological variation

We next tested the reproducibility of hcHCR by performing dHCR for *TNF* and qHCR for *IL1B* in human primary monocytes obtained from 3 unrelated healthy donors. For each biological replicate, the assays were performed in 8 technical replicates. To incorporate measurements of up- or down-regulation of gene expression in this assessment, we performed *in vitro* stimulation with LPS for 30 minutes, followed by treatment with methylprednisolone or vehicle. We observed a 14.6-fold induction of *TNF* expression when comparing untreated with LPS-stimulated monocytes (Fig 5A). As expected, *in vitro* treatment of LPS-treated cells with MP reduced *TNF* expression by half when compared with the LPS-stimulated samples that were not treated with MP (Fig. 5A). Similarly, we observed a 26.4-fold induction of *IL1B* in the presence of LPS, and a reduction of *IL1B* expression by half when compared to the LPS-stimulated samples when monocytes were subsequently treated with MP (Fig. 5B). These results were consistent in the 3 biological replicates (Fig. 5A and 5B).

**Figure 5.**
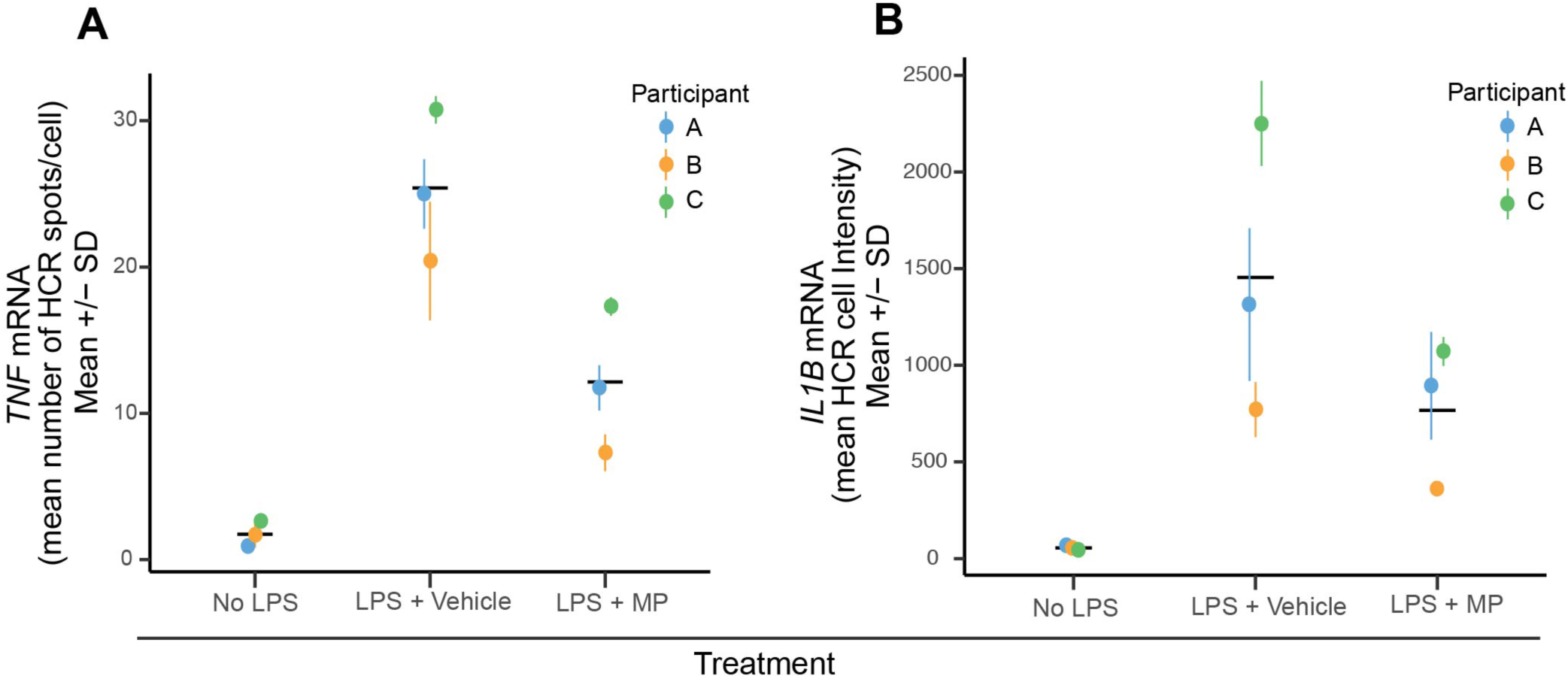
Assessment of technical and biological variation in hcHCR data. Primary human monocytes were obtained from 3 unrelated healthy donors and dHCR was performed with *TNF* as the target mRNA (A) and qHCR was performed with *IL1B* as the target mRNA (B). Measurements were performed at baseline (no LPS), after stimulation with 1 ng/mL LPS for 30 minutes followed by vehicle treatment (0.1% EtOH) for 2 hours (LPS + Vehicle), or after stimulation with 1 ng/mL LPS for 30 minutes followed by methylprednisolone treatment (200 µg/dL) for 2-hours (LPS + MP). Each dot represents one biological replicate. Colored error bars display the mean ± SD of 8 technical replicates measured in each biological replicate. Black error bars display the mean ± SD of the biological replicates.

We then assessed the level of technical variation in the assay. For dHCR with *TNF* as the target mRNA, the coefficient of variation (CV) for the 8 technical replicates (averaged across the 3 biological replicates) had a mean of 29.29% at baseline, 10.77% after LPS stimulation, and 11.34% in cells stimulated with LPS then treated with MP (Fig. 5A). For qHCR with *IL1B* as the target mRNA, the CV had a mean of 14.01% at baseline, 19.4% after LPS stimulation, and 16.08% in cells stimulated with LPS then treated with MP (Fig. 5B).

Finally, we evaluated the level of variation across biological replicates for either gene across conditions. We found evidence of substantial biological variation in the expression of both genes at baseline and in response to stimulation and treatment (Fig. 5A and 5B). At baseline, the mean *TNF* spots/cell was 1.74 with SD = 0.86 and CV = 49.45%. After LPS stimulation, the mean was 25.38 with SD = 5.18 and CV = 20.41%. In cells stimulated with LPS then treated with MP, the mean was 12.12 with SD = 5.01 and CV = 41.38%. For *IL1B* expression, the mean cell intensity at baseline was 54.68 with SD = 11.92 and CV = 21.81%. After LPS stimulation, the mean was 1445.15 with SD = 749.04 and CV = 51.83%. In cells stimulated with LPS then treated with MP, the mean was 775.07 with SD = 370.09 and CV = 47.75%. These results highlight the importance of considering both technical and biological sources of variation when working with primary human cells, and also highlight the ability of the hcHCR to account for both, given the relatively small numbers of cells per well that are required.

## Materials and Methods

### Cell purification

Human peripheral blood hematopoietic cells were obtained from the Department of Transfusion Medicine at the National Institutes of Health (NIH) Clinical Center, under NIH study 99-CC-0168, Collection and Distribution of Blood Components from Healthy Donors for In Vitro Research Use, which was approved by the Clinical Center’s Institutional Review Board. Peripheral blood was collected in vacutainer EDTA tubes (Becton Dickinson; cat. no. 366643) for all the participants. Mononuclear cell subsets were obtained by isolation of PBMCs by gradient centrifugation in SepMate tubes (STEMCELL Technologies; cat. no. 85460), with Ficoll-Paque PLUS (GE Healthcare Life Sciences; cat. no. 17-1440-03). Immediately after isolation, and before treatment, mononuclear cells were incubated overnight in RPMI 1640 (ThermoFisher Scientific, cat. no. 11875093) and 10% FBS at 4°C, followed by immunomagnetic enrichment for the specific cell subset with EasySep Human cell enrichment kits (STEMCELL Technologies). B lymphocytes and CD4^+^ T lymphocytes were isolated from PBMCs by negative selection (STEMCELL Technologies; cat. nos. 19054 and 19052, respectively). Monocytes were isolated from PBMCs by positive selection (STEMCELL Technologies; cat. no. 17858), to ensure inclusion of the CD14^+^/CD16^+^ fraction, which would be excluded with the use of a negative-selection kit. Neutrophils were freshly isolated directly from whole blood by negative selection immunomagnetic purification with the EasySep Direct Human Neutrophil Isolation Kit (STEMCELL Technologies; cat. no. 19666).

### Cell adherence to multiple substrates

Custom EvaluPlate Attachment Surfaces plates (BioMedTech Laboratories, Inc.) were generated by coating wells of CellCarrier-384 Ultra imaging microplates (PerkinElmer, cat. no. 6057300) with 8 different cell culture matrices: MS-1, MS-2, MS-3, 3D-Hydrogel, Poly-D-Lysine (PDL), Synthetic Poly-Amine (SPA), 3D-Hydrogel PDL, 3D-Hydrogel SPA. 384-well imaging coated plates were stored at - 20°C. They were equilibrated to room temperature for an hour after removal from storage. Each of the four cell types, B cells, monocytes, neutrophils and CD4+ T cells from three unrelated healthy participants were plated at three different concentrations of 100000, 50000 and 25000 cells/well in 3 technical replicates. All the cells were plated in 40 µL RPMI 1640 and 10% FBS. After incubation for 2 to 4 hours at 37°C, 5% CO_2_, cells were fixed by adding the fixative directly to the medium to a final concentration of 4% paraformaldehyde (PFA) (Electron Microscopy Sciences; cat. no. 15714) using the automated liquid handler BlueWasher (BLUE CAT BIO). The plates were incubated for 15-min at room temperature followed by three washes with 1X PBS and by permeabilization with 70% ethanol at -20°C for 20 to 70 hours. Before imaging, cells were washed multiple times with 1x PBS to mimic the HCR protocol washes, stained with DAPI (2.5 ng/µL) per well for 20-min at room temperature. After discarding DAPI, 50 µL of 1x PBS was added to each well and imaged as described further below.

### LPS titration for *TNF* and *IL1B* mRNA expression

Monocytes from three unrelated healthy participants were plated in 30 µL of RPMI 1640 and 10% FBS on CellCarrier-384 well Ultra Microplates coated with PDL (PerkinElmer; cat. no. 6057500) at a concentration of 100,000 cells/well. After 2 hours of rest at 37°C, 5% CO_2_, cells were stimulated in technical replicates with six different LPS concentrations (0, 0.01, 0.1, 1, 10 and 100 ng/mL) at five time points (0, 15, 30, 60 and 120-min). Negative (LPS 0 ng/mL) and positive (LPS 100 ng/mL) 120-min stimulation hairpin controls were also included (two hairpins + *TNF* probe, two hairpins + *IL1B* probe, two hairpins without primary probe). All the wells were fixed, washed and permeabilized overnight as described in the Cell adherence to multiple substrate section. Followed by RNA-HCR.

### RNA Hybridization Chain Reaction (HCR)

For pre-hybridization, the 70% ethanol permeabilization buffer was discarded by inverting the plate and pat drying, followed by aspiration of the remaining buffer with a sterile 200 µL tip attached to a sterile 2 mL aspirating pipette. Plates were air dried for 10-min at room temperature to remove any residual ethanol. Wells were rehydrated by three 5-min washes in 80 µL of 5X Saline-Sodium Citrate, 0.1% Tween20 (5X SSC-T) buffer (Ambion, cat. no. AM9763) at room temperature. HCR probe sets, amplifiers, probe hybridization buffer, amplifier buffer (Molecular Instruments, v3.0) were used for HCR. For equilibration, 16 µL of probe hybridization buffer was added to each well and incubated for 10 min at 37°C. Primary probe-hybridization mix was prepared by multiplexing pairs (odd and even) of each probe set to a final concentration of 2 nM in probe hybridization buffer. After aspirating the equilibration buffer from each well, 11 µL of pre-warmed (37°C) probe-hybridization mix was added to each well. All the plates were immediately sealed with an aluminum seal (Fisher Scientific, cat. no. 07-000-379) and incubated in a humidified 37°C incubator for 12-18 hours. After overnight incubation, the probe-hybridization mix was aspirated followed by four 15 min washes in a 37°C water bath with pre-warmed solutions (37°C) to remove the excess probes. The first wash was performed with 75% of probe wash buffer/ 25% 5X SSC-T, followed by a wash with 50% of probe wash buffer/ 50% 5X SSC-T, then by a wash with 25% of probe wash buffer/ 75% 5X SSC-T, and finally by a wash with 100% 5X SSC-T. After the fourth wash, one more wash was performed with 5X SSC-T for 5 min at RT. To equilibrate the plates before HCR amplification, the wash buffer from each well was aspirated and 16 µL of amplification buffer was added and incubated at room temperature for 30 min. Amplification hairpins were prepared by thawing a pair of hairpins (H1 and H2, either labelled with Alexa488 or Alexa647) on ice for each probe used, followed by snap heating at 95°C for 1 min 30 sec and equilibrating to room temperature for 30 min. Each hairpin was multiplexed to a final concentration of 60 nM in amplification buffer (2X SSC, 0.1% Triton-X 100, 10% dextran sulfate). Before adding the multiplexed hairpins to the wells, the amplification buffer was aspirated and 11 µL of multiplexed hairpin mix was added. Plates were incubated at RT for 45-min, followed by two 30 min and one 5 min wash with 5X SSC-T at RT. Cells were stained for DAPI (2.5 ng/µL) per well for 20 min at room temperature and either imaged right away, or stored at 4°C in 50 µL of 1X PBS.

### High-throughput image acquisition

Plates stained with HCR probe sets and amplification hairpins were imaged on a CV7000S high-throughput spinning disk confocal microscope (Yokogawa, Japan). Samples were first excited using a 405 nm solid state laser, a 405/488/561/640 nm excitation dichroic mirror, a 60X water objective (NA 1.2), a 568-emission dichroic mirror, and a 445/45 nm bandpass emission filter to detect the DAPI signal. The second exposure used simultaneous excitation with 488 and 640 nm excitation lasers, the same excitation dichroic mirror, objective, and emission dichroic mirror as the first exposure, and 525/50 nm and 676/29 nm bandpass emission filters to detect the Alexa488 and Alexa647, respectively. For both exposures, we used 2 sCMOS cameras (2560 × 2160 pixels) with bin setting of 2×2 to acquire 3D z-stacks of 5 images at 1 μm intervals, which were maximally projected, background and shading corrected, and registered on the fly during image acquisition by Yokogawa proprietary algorithms, and saved as .tif files. For each well, we acquired 4 fields of view (FOV).

### High-content image analysis

Images were imported and analyzed in Columbus 2.7 or 2.8 (PerkinElmer, Waltham, MA). Briefly, nuclei were segmented using the DAPI channel, and dilated by a fix percentage to generate an approximate cell body region of interest. For digital HCR (dHCR, Choi, 2018) HCR foci were first detected over the cell body region using Columbus spot finding algorithm C, and then filtered using a user-trained Fisher Linear Discriminant classifier based on fluorescence intensity and contrast. The output of dHCR was number of HCR spots per cell. For quantitative HCR (qHCR, Choi et al. 2018) the mean fluorescence intensity in the HCR channel was measured over the cell body region and used as the output measurement. Single cell results were exported from Columbus as text files.

### Data display

Single-cell results generated in Columbus were used to generate .pdf plots in R (3.6.3, R Core Team) and RStudio Desktop (RStudio). Original .tif files were processed in FIJI/ImageJ (NIH) by changing only brightness and contrast settings over the entire FOV and by maintaining them constant among different experimental conditions in the same figure panel. Grayscale 16-bit images from different channels were merged, and then converted to 8-bit RGB format.

For visualization of image data with standard flow cytometry software, single-cell results were exported from Columbus in comma-separated-value (CSV) format. The files were then converted to Flow Cytometry Standard (FCS) format with the CsvToFcs module of GenePattern (Spidlen et al. 2013). Data in FCS format were displayed as bivariate dot plots with FlowJo, v10.

Plots and images were assembled into figures with Adobe Illustrator (Adobe).

### Statistical analysis

To assess the difference in cell attachment across substrates, a linear mixed-effects model was fitted to account for the repeated measurements on cells obtained from the same participants and cultured on wells coated with different substrates. To assess the difference of retention ratios across cell concentrations, a linear mixed-effects model was fitted to account for the repeated measurements on cells obtained from the same participants and cultured at different cell concentrations. In both cases, the resulting p-values reflect the significance of the within-participant differences under different conditions.

### LPS stimulation and methylprednisolone treatment

Monocytes from three unrelated healthy participants were plated as described in the LPS titration for *TNF* and *IL1B* mRNA expression section. After 2-hours of rest at 37°C, 5% CO_2_, cells either received no LPS or were exposed to two conditions, LPS (1 ng/mL) stimulation for 30-min followed by vehicle (0.1% EtOH) or methylprednisolone (200 µg/dL) (Millipore Sigma, cat. no. M0639) treatment for 2 hours. Cells were fixed, washed and permeabilized overnight as described in the Cell adherence to multiple substrate section. This was followed by RNA-HCR.

## Discussion

We have developed a method for HCI quantification of transcript abundance at the single-cell level in primary human immune cells. The method combines the high sensitivity of HCR for the quantification of gene expression with the speed, scalability, and technical reproducibility of high-content imaging, and we abbreviate it as hcHCR. While hcHCR is not intended to replace other methods for medium-or high-throughput quantification of gene expression, it has advantages that make it especially suitable for a range of applications.

Compared to scRNA-seq, hcHCR is considerably simpler and less expensive, making it more scalable. Because cells are imaged directly on the plate and not captured, it also has the potential to reduce biases in cell representation introduced by the capture step. In addition, genes with very low transcript abundance are likely to be excluded in scRNA-seq, yet they are easily visualized and quantified by hcHCR. Genes with high transcript abundance in very few cells can also be excluded at the capture step, or lead to misclassification of the cell in scRNA-seq pipelines; by simply imaging more fields of view, hcHCR can be scaled up to quantify 10,000 – 20,000 cells per well, thus increasing the likelihood of accurate quantification of such genes. On the other hand, scRNA-seq has the advantage of simultaneously assaying up to a few thousand transcripts on any given cell which, although a shallow representation of the transcriptome, is much broader than what can be achieved by hcHCR. Bulk RNA-seq can offer a deeper analysis of the entire transcriptome, at the expense of single-cell resolution. We see these methods as complementary, with RNA-seq offering the possibility of identifying a subset of genes that could be appropriate markers for the response to a specific perturbation, and hcHCR providing a scalable way to assay the expression of these marker genes in a wide range of conditions, doses, and cell types.

Real-time quantitative PCR (qPCR) is another technique for quantification of transcript abundance. The most scalable and most widely used form of qPCR involves bulk measurements of RNA obtained from all the cells in a well. The obvious advantage of hcHCR is the single-cell resolution, which allows the quantitative evaluation of subsets of cells within a well that may be more or less transcriptionally active or responsive to perturbation. With the ability to multiplex, hcHCR also offers the advantage of assessing correlation in the magnitude of the response at the level of different genes in individual cells. For example, in our validation studies it was evident that the human monocytes that responded to LPS with the strongest induction of *TNF* expression were the same cells that responded with the strongest induction of *IL1B* expression. Methods for single-cell qPCR have been developed. Like scRNA-seq, these rely on a capture step, which reduces their scalability, increases cost, and introduces the possibility of capture biases compared to imaging of the cells directly on the plate.

HCR has been successfully applied to flow cytometry (Choi et al. 2018), and that method offers many of the advantages of hcHCR, including high sensitivity and single-cell resolution. Cost and scalability are advantages of hcHCR over HCR-flow cytometry. Conversely, there are specific situations in which HCR-flow cytometry may be preferable, for example in cases where cell viability must be preserved after measurement for downstream applications.

Considering its relative advantages and limitations, we believe that hcHCR will be most suitable and cost-effective for medium-throughput screens for the biological effects of perturbing agents such as chemical compounds, RNAi, or targeted genome editing. The applicability of the method to primary cells is important, as it allows for screens with multiple biological replicates and opens the possibility of studying cells directly obtained from patients with the particular disease state being studied.

## Acknowledgements

This study was funded by the Intramural Research Programs of the National Institute of Allergy and Infectious Diseases; the National Cancer Institute, and the National Institute of Arthritis and Musculoskeletal and Skin Diseases, all at the National Institutes of Health.

## Bibliography

Cannarile L, Zollo O, D’Adamio F, Ayroldi E, Marchetti C, Tabilio A, Bruscoli S, Riccardi C. 2001. Cloning, chromosomal assignment and tissue distribution of human GILZ, a glucocorticoid hormone-induced gene. Cell Death Differ 8: 201–203.

Chen AR, McKinnon KP, Koren HS. 1985. Lipopolysaccharide (LPS) stimulates fresh human monocytes to lyse actinomycin D-treated WEHI-164 target cells via increased secretion of a monokine similar to tumor necrosis factor. J Immunol 135: 3978–3987.

Chen G, Ning B, Shi T. 2019. Single-Cell RNA-Seq Technologies and Related Computational Data Analysis. Front Genet 10: 317.

Choi HMT, Chang JY, Trinh LA, Padilla JE, Fraser SE, Pierce NA. 2010. Programmable in situ amplification for multiplexed imaging of mRNA expression. Nat Biotechnol 28: 1208–1212.

Choi HMT, Schwarzkopf M, Fornace ME, Acharya A, Artavanis G, Stegmaier J, Cunha A, Pierce NA. 2018. Third-generation in situ hybridization chain reaction: multiplexed, quantitative, sensitive, versatile, robust. Development 145.

D’Adamio F, Zollo O, Moraca R, Ayroldi E, Bruscoli S, Bartoli A, Cannarile L, Migliorati G, Riccardi C. 1997. A new dexamethasone-induced gene of the leucine zipper family protects T lymphocytes from TCR/CD3-activated cell death. Immunity 7: 803–812.

Esner M, Meyenhofer F, Bickle M. 2018. Live-Cell High Content Screening in Drug Development. Methods Mol Biol 1683: 149–164.

Franco LM, Gadkari M, Howe KN, Sun J, Kardava L, Kumar P, Kumari S, Hu Z, Fraser IDC, Moir S, et al. 2019. Immune regulation by glucocorticoids can be linked to cell type-dependent transcriptional responses. J Exp Med 216: 384–406.

Gibellini L, De Biasi S, Porta C, Lo Tartaro D, Depenni R, Pellacani G, Sabbatini R, Cossarizza A. 2020. Single-Cell Approaches to Profile the Response to Immune Checkpoint Inhibitors. Front Immunol 11: 490.

Gioia L, Siddique A, Head SR, Salomon DR, Su AI. 2018. A genome-wide survey of mutations in the Jurkat cell line. BMC Genomics 19: 334.

Hauser H. 2015. Cell Line Development. In Animal Cell Culture (ed. M. Al-Rubeai), Vol. 9 of Cell Engineering, pp. 1–25, Springer International Publishing, Cham.

Hodge S, Hodge G, Flower R, Han P. 1999. Methyl-prednisolone up-regulates monocyte interleukin-10 production in stimulated whole blood. Scand J Immunol 49: 548–553.

Hughes JP, Rees S, Kalindjian SB, Philpott KL. 2011. Principles of early drug discovery. Br J Pharmacol 162: 1239–1249.

Hwang B, Lee JH, Bang D. 2018. Single-cell RNA sequencing technologies and bioinformatics pipelines. Exp Mol Med 50: 96.

Kornbluth RS, Edgington TS. 1986. Tumor necrosis factor production by human monocytes is a regulated event: induction of TNF-alpha-mediated cellular cytotoxicity by endotoxin. J Immunol 137: 2585–2591.

Lavrentieva A. 2018. Essentials in cell culture. In Cell Culture Technology (eds. C. Kasper, V. Charwat, and A. Lavrentieva), Learning materials in biosciences, pp. 23–48, Springer International Publishing, Cham.

Mittelman D, Wilson JH. 2013. The fractured genome of HeLa cells. Genome Biol 14: 111.

Pegoraro G, Misteli T. 2017. High-Throughput Imaging for the Discovery of Cellular Mechanisms of Disease. Trends Genet 33: 604–615.

Pichon X, Lagha M, Mueller F, Bertrand E. 2018. A growing toolbox to image gene expression in single cells: sensitive approaches for demanding challenges. Mol Cell 71: 468–480.

Pope SD, Medzhitov R. 2018. Emerging principles of gene expression programs and their regulation. Mol Cell 71: 389–397.

Querido E, Dekakra-Bellili L, Chartrand P. 2017. RNA fluorescence in situ hybridization for high-content screening. Methods 126: 149–155.

Spidlen J, Barsky A, Breuer K, Carr P, Nazaire M-D, Hill BA, Qian Y, Liefeld T, Reich M, Mesirov JP, et al. 2013. GenePattern flow cytometry suite. Source Code Biol Med 8: 14.

Stubbington MJT, Rozenblatt-Rosen O, Regev A, Teichmann SA. 2017. Single-cell transcriptomics to explore the immune system in health and disease. Science 358: 58–63.

Sun J, Li N, Oh K-S, Dutta B, Vayttaden SJ, Lin B, Ebert TS, De Nardo D, Davis J, Bagirzadeh R, et al. 2016. Comprehensive RNAi-based screening of human and mouse TLR pathways identifies species-specific preferences in signaling protein use. Sci Signal 9: ra3.

Tiligada E, Ishii M, Riccardi C, Spedding M, Simon H-U, Teixeira MM, Cuervo MLC, Holgate ST, Levi-Schaffer F. 2015. The expanding role of immunopharmacology: IUPHAR Review 16. Br J Pharmacol 172: 4217–4227.

Waage A, Bakke O. 1988. Glucocorticoids suppress the production of tumour necrosis factor by lipopolysaccharide-stimulated human monocytes. Immunology 63: 299–302.

Zhou B, Ho SS, Greer SU, Spies N, Bell JM, Zhang X, Zhu X, Arthur JG, Byeon S, Pattni R, et al. 2019. Haplotype-resolved and integrated genome analysis of the cancer cell line HepG2. Nucleic Acids Res 47: 3846–3861.

